# Analysis of Categorical Data with Logistic Regression and the Cochran–Mantel–Haenszel Tests in Biological Experiments

**DOI:** 10.1101/2024.11.14.623695

**Authors:** Rebecca Androwski, Tatiana Popovitchenko, Joelle Smart, Sho Ogino, Guoqiang Wang, Christopher Rongo, Monica Driscoll, Jason Roy

**Author notes:** Co-first authors (Rebecca Androwski, Tatiana Popovitchenko).

## Abstract

The choice of statistical test is a fundamentally important one when analyzing experimental data. Here, we consider the question of categorical data, defined by their properties (for example color) rather than by continuous numbering. Using simple and complex example datasets generated from *Caenorhabditis elegans* research, we conduct a statistical analysis of (1) a rare cellular event involving the formation of a neuronal extrusion called an exopher, and of (2) a variable behavioral response across a timescale. Two tests we use here are the Cochran– Mantel–Haenszel (CMH) test and logistic regression. These two tests pose practical challenges to researchers that include lack of easy access to statistical software and the need for prior programming knowledge. To this end we provide step-by-step tutorials and example code. We emphasize the flexibility of logistic regression in handling both simple and complex datasets, emphasizing the capacity of logistic regression to provide more comprehensive insights into experimental outcomes than simpler tests like CMH. By analyzing real biological examples and demonstrating their analysis with R code, we provide a practical guide for biologists to enhance the rigor and reproducibility of categorical data analysis in experimental studies.

## Introduction

Genetic research has historically leveraged the advantages of large population sizes and, for some model systems, clonal reproduction to examine biological principles in action. Discoveries from genetics studies have led to revolutionary breakthroughs in modern science and thanks to large population sizes, infrequent, yet biologically significant, phenomena can be detected and quantified in genetic models. The significance of these infrequent events can be difficult to represent numerically, as occasional events are often outnumbered by prevalent baseline biological outcomes. Appropriate statistical analysis of categorical data provides a layer of confidence and rigor in its interpretation, yet deciding on an appropriate statistical test of categorical data can be challenging. Here, we consider the challenge of detecting statistical differences between unbalanced groups, particularly those that are measured categorically.

Categorical outcomes are common in both cell biological experiments and behavioral assays involving binomial or multinomial datasets. Categorical data (or categorical variables) are data that can be described by a specific classification. Such data can include nominal categorical variables, which do not have a natural specific order (e.g., color or genotype); ordinal variables, which are ranked categories (e.g., level of damage or progression of disease); and dichotomous variables, which are variables with only two outcomes (e.g., alive or dead). Dichotomous variables are abundant in the *Caenorhabditis elegans* (*C. elegans*) experimental repertoire (e.g., alive or dead, on food or off food, moving or not moving). In one of the examples we detail here, we focus on neuronal production of large extracellular vesicles (called exophers) under conditions of proteostress^1^. Exophers bud directly from the neuronal cell body (commonly one per neuron) and can be equal in size to the cell body itself (∼5µm). In this case, individual animals can be easily categorized by the presence or absence of the exopher and the frequency of exopher events can be assessed over a large sample size of nematodes.

Regarding data analysis, the dichotomous variables are whether the neuron produced an exopher or did not produce an exopher such that an exopher frequency (number of exophers observed/total animals assayed) can be calculated for a sample population.

Multiple statistical approaches can be used to analyze categorical variables. Categorical data is often plotted as a percentage (%) of the whole, also known as a frequency. Since percentage/frequency scores appear to be continuous data, investigators might incorrectly use the *t-*test or one-way ANOVA to analyze these categorical data. The frequency of a binary event from a single trial would more appropriately be analyzed using a Chi-square or Fisher’s exact test. When comparing replicate trials of categorical outcomes, a better option is the Cochran– Mantel–Haenszel test (CMH)^2^. CMH tests the association between a binary independent variable, commonly called the predictor in statistics, and a binary dependent variable, called the outcome. CMH uses outcomes grouped into categories for analysis, summarizes those categories into contingency tables, and calculates an odds ratio (a likelihood that an outcome is linked to a variable). However, the CMH method is limited to situations where there is a binary outcome and a binary treatment, such that the data fall into relatively few categories. Further, the CMH test requires the assumption of a common odds ratio between outcomes (e.g., the odds ratio between treatment and outcome is the same in both categories).

Another statistical approach to analyzing categorical data is logistic regression. Logistic regression is a version of linear regression that examines the relationship between an independent variable/predictor and its outcome^3^. A logistic regression models the log-odds of the outcome and can be depicted as an S-shaped curve on a plane. Yes/no values can be represented as one for yes and zero for no, and the slope of the curve between the top and bottom will indicate how likely an animal will be at one or zero. Logistic regression can be used for categorical outcomes, can include binary (i.e. alive or dead) or continuous predictors (i.e. increasing doses of a drug), and can include variables to control for factors such as replicate trials or time. Thus, logistic regression can be tailored to the specific experimental design and offers the capacity to analyze data over multiple variables^4^.

Logistic regression is a powerful and adaptable test but is not commonly taught in introductory statistics classes. For this reason, logistic regression analysis may be underutilized in the field. Here, we emphasize that the less commonly applied logistic regression model provides the most comprehensive statistical analysis for categorical data. To illustrate the application of this statistical test, we apply CMH and logistic regression to an example of large vesicle extrusion from neurons. We also apply an ANOVA analysis and logistic regression to locomotory behavior in multiple genetic backgrounds over a time course. We include advice for improved data hygiene, a basic R Studio tutorial provided in the supplement, and sample R code for direct implementation (found at https://github.com/Randrowski/Logistic-Regression-for-Biologists.git).

### Logistic Regression for a simple example problem

Our first example dataset for how to start using logistic regression deals with the quantification of exophers in *C. elegans* neurons. Large vesicles can be extruded from proteo-stressed touch sensory neurons in *C. elegans* via a process called exophergenesis^5^ (Figure 1A). Exopher events at a specific neuron are rare for each animal; for example, only 5-20% of wild-type animals typically produce exophers from their ALM neuron. In our exopher dataset the categorical variable is the presence of the exopher itself: exopher or no exopher (Figure 1A, B). While the exopher event is rare, we can easily analyze large populations (>50 animals) and perform >3 biological replicates and multiple trials. Thus, while enough observations can be made for a conclusion^6^ the resulting data are severely unbalanced and analysis is complicated^7^. Statistical tests, such as t-tests that rely on a comparison of means or ANOVAs that rely on a sum of squares, can dismiss exopher production entirely. Yet, as we have argued, exopher production is a consequential biological event that merits analysis^5^. We required a statistical test that could correctly assess categorical results across replicate trials, would allow for the high variation in baseline that is inherent to our biological model, and could identify differences among relatively infrequent events where total sample sizes could vary amongst trials.

**Figure 1.**
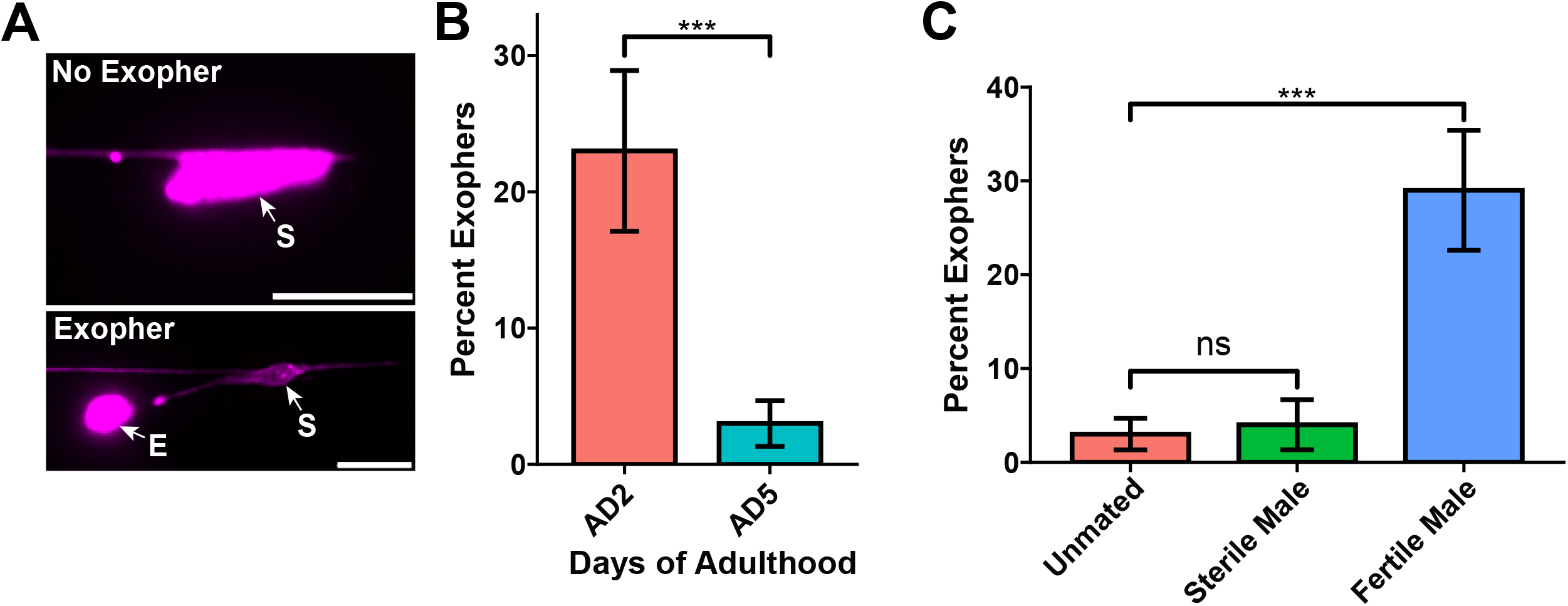
Analysis of exophers and animal movement using logistic regression. **A)** Fluorescence micrographs show exophers “E” are extruded from the neuronal soma “S” and are scored as a binary event, as exopher or no exopher. The touch receptor neuron, ALM is labeled with *Pmec-4m::Cherry* to allow for easy assessment of cell soma and exopher. Scale bar, 10 µm. **B)** Exopher frequency decreases on adult day 5 (AD5) compared to the frequency on adult day 2 (AD2). *** p<0.001 by both the Cochran–Mantel Haenszel (CMH) test and logistic regression, N = 50 per trial over two trials. **C)** Comparison of multiple mating schemes and their effect on exopher frequency. On AD5, there is no difference between unmated and animals mated with sterile males while animals mated with fertile males results in a significant increase in exopher frequency. *** p<0.001, “ns” = not significant by logistic regression, N = 50 per trial over two trials. Error bars = standard error of the percentage. All example exopher data were originally published in Wang et al 2024.

### Comparison of CMH vs. logistic regression

We found that the CMH statistical test met the criteria for analysis of exopher frequencies when single comparisons were made. For example, we compared exopher frequency in young adults (adult day 2) to that in older adults (adult day 5) in replicate trials; CMH analysis yielded a P value of p= 0.0000624 (Figure 1B). Evaluation of the same data with the logistic regression model yielded a similar significance level (p=0.000301). A comparison of features between the CMH and logistic regression analyses is summarized in Table 1. In this work, we found that a specific formatting of collected data enabled easy application of the statistical test. Step-by-step instructions for reformatting data for CMH, including annotated PDFs for both simple and multiple comparisons, can be found in our Github repository (https://github.com/Randrowski/Logistic-Regression-for-Biologists.git). As an alternative to using R, we also provide an excel spreadsheet that will calculate a CMH p-value according to the *Handbook of Biological Statistics*^8^. The CMH statistic can also be applied through the following web application, https://biostats-ashen.vercel.app/.

**Table 1.**
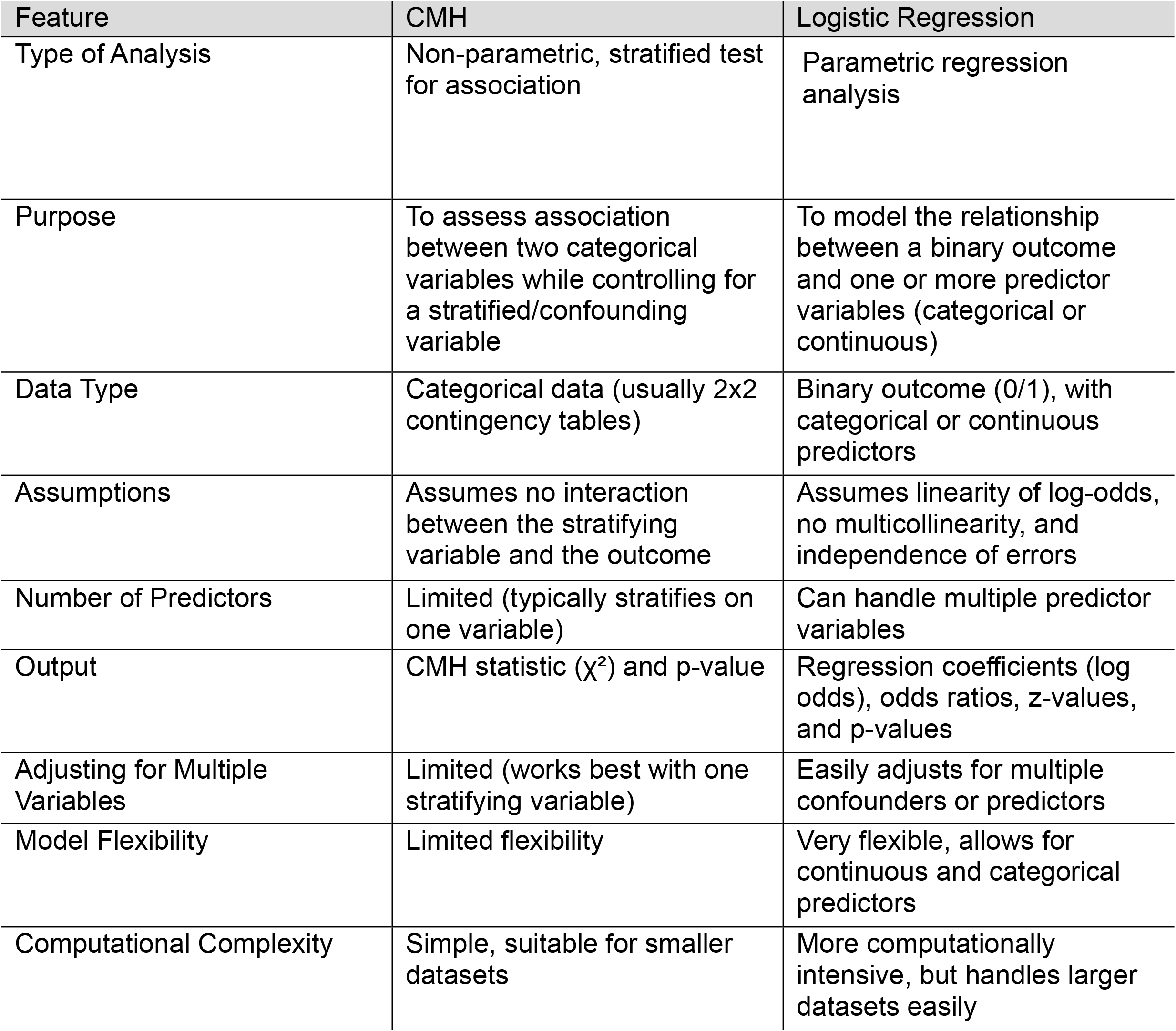
Comparison of features between CMH and Logistic Regression. The CMH test is simpler and more specialized, allowing for analysis of categorical data that has one variable and controls for stratification (dividing into groups for analysis) – meaning it adjusts for subgroup differences, ensuring that the association between the independent (predictor) and dependent (outcome) variables is evaluated within each stratum separately. This control allows for a more accurate analysis by reducing bias due to confounding factors that might vary across the strata. Logistic regression is more powerful and versatile, allowing for the modeling of multiple predictors. However logistic regression relies on stronger assumptions (that mean and variance are a normal distribution) and requires more computational effort.

| Feature | CMH | Logistic Regression |
| --- | --- | --- |
| Type of Analysis | Non-parametric, stratified test for association | Parametric regression analysis |
| Purpose | To assess association between two categorical variables while controlling for a stratified/confounding variable | To model the relationship between a binary outcome and one or more predictor variables (categorical or continuous) |
| Data Type | Categorical data (usually 2x2 contingency tables) | Binary outcome (0/1), with categorical or continuous predictors |
| Assumptions | Assumes no interaction between the stratifying variable and the outcome | Assumes linearity of log-odds, no multicollinearity, and independence of errors |
| Number of Predictors | Limited (typically stratifies on one variable) | Can handle multiple predictor variables |
| Output | CMH statistic ($\chi^2$) and p-value | Regression coefficients (log odds), odds ratios, z-values, and p-values |
| Adjusting for Multiple Variables | Limited (works best with one stratifying variable) | Easily adjusts for multiple confounders or predictors |
| Model Flexibility | Limited flexibility | Very flexible, allows for continuous and categorical predictors |
| Computational Complexity | Simple, suitable for smaller datasets | More computationally intensive, but handles larger datasets easily |

CMH and logistic regression both report an odds ratio. The odds ratio is a likelihood statistic and compares the likelihood of something happening in one group compared to it happening in another group. While probability is the likelihood of something happening out of the total, the odds ratio compares the odds between two groups and is especially informative in experiments in which there is a control condition. An odds ratio of “1” indicates that an event is equally likely in both groups while an odds ratio of “2” indicates that something is twice as likely to occur in one group compared to the other. Additionally, one can also report the 95% confidence interval for the odds ratio, which provides a range of values within the 95% confidence interval in which the true odds ratio lies and gives some insight into the precision of the estimated odds ratio.

Statistical significance can be supported from the confidence interval of the odds ratio by simply determining if the confidence interval overlaps with “1”. Since an odds ratio of “1” indicates no difference between the treatments, an overlap of the 95% confidence interval with “1” would suggest the difference is not significant, whereas a confidence interval that does not overlap “1” suggests that there is a significant difference between the treatments (annotated code for calculating the odds ratio and 95% confidence intervals are included in https://github.com/Randrowski/Logistic-Regression-for-Biologists.git).

Notably, logistic regression supplies more descriptive information in this simple comparison by reporting the direction of the change in the form of a z-value. The z-value describes how far the data lies from the average of the control dataset. An increase in frequency for the experimental versus the control would yield a positive numerical value for the z statistic; a negative z statistic would indicate a decrease in frequency. In our exopher example, we tested whether day 5 adults have a higher or lower frequency of exophergenesis compared to day 2 controls. The z-value was z=-3.614, indicating that the exopher frequency in day 5 adults was lower than that in day 2 adults. The z-value also provides an effect size, representing standard deviations from the mean of the control data. Staying with our exopher example, the magnitude of the z-value (z=-3.614) indicates that the frequency of exophers was more than three and a half standard deviations below that of the younger animal control. Thus, logistic regression analysis demonstrates that *C. elegans* hermaphrodites experience a large and statistically significant decrease in exopher frequency by adult day 5. CMH and logistic regression thus both provide a P statistic and an odds ratio, but logistic regression further provides information on the direction of the change as well as the relationship to the mean; this leads us to choose the logistic regression test in the analysis of exophergenesis.

### Data collection framework

An important feature of a rigorous and reproducible statistical analysis is in the formatting of the initial data sheet or data frame. This careful formatting is otherwise known as data hygiene. In general, a single individual assessed for a phenotype should occupy a single row in the data sheet, whereas features of all individuals should be organized into columns. The simplest arrangement would be two columns labeled genotype and outcome, indicated by a one or a zero. However, there could be other additional relevant information that needs to be considered in the comparison. For example, the anoxia-behavioral dataset example we consider below contains four columns (trial, time, genotype, and outcome), allowing one to assess a dichotomous outcome (movement vs no movement in our example) along a time series and across trials.

Importantly, attention to careful data hygiene ensures that data are formatted for import into R scripts. As an added benefit, standardized formatting of data allows for easy sharing within lab groups and with biostatistical collaborators (Figure 2). It may seem more useful to have data sheets formatted with color coded text or columns, which we often see in biological research, but in practice the documentation of an observation on an individual level leads to enhanced accessibility, rigor, reproducibility, and application. We present sample data collection spreadsheets, R code for reformatting data for analysis, and an annotated PDF of the code (highlighting terms that might be specific to an individual experiment) in Extended Data sections for each of the datasets discussed here, thereby demonstrating how we efficiently process data from collection into an analysis-ready dataset.

**Figure 2.**
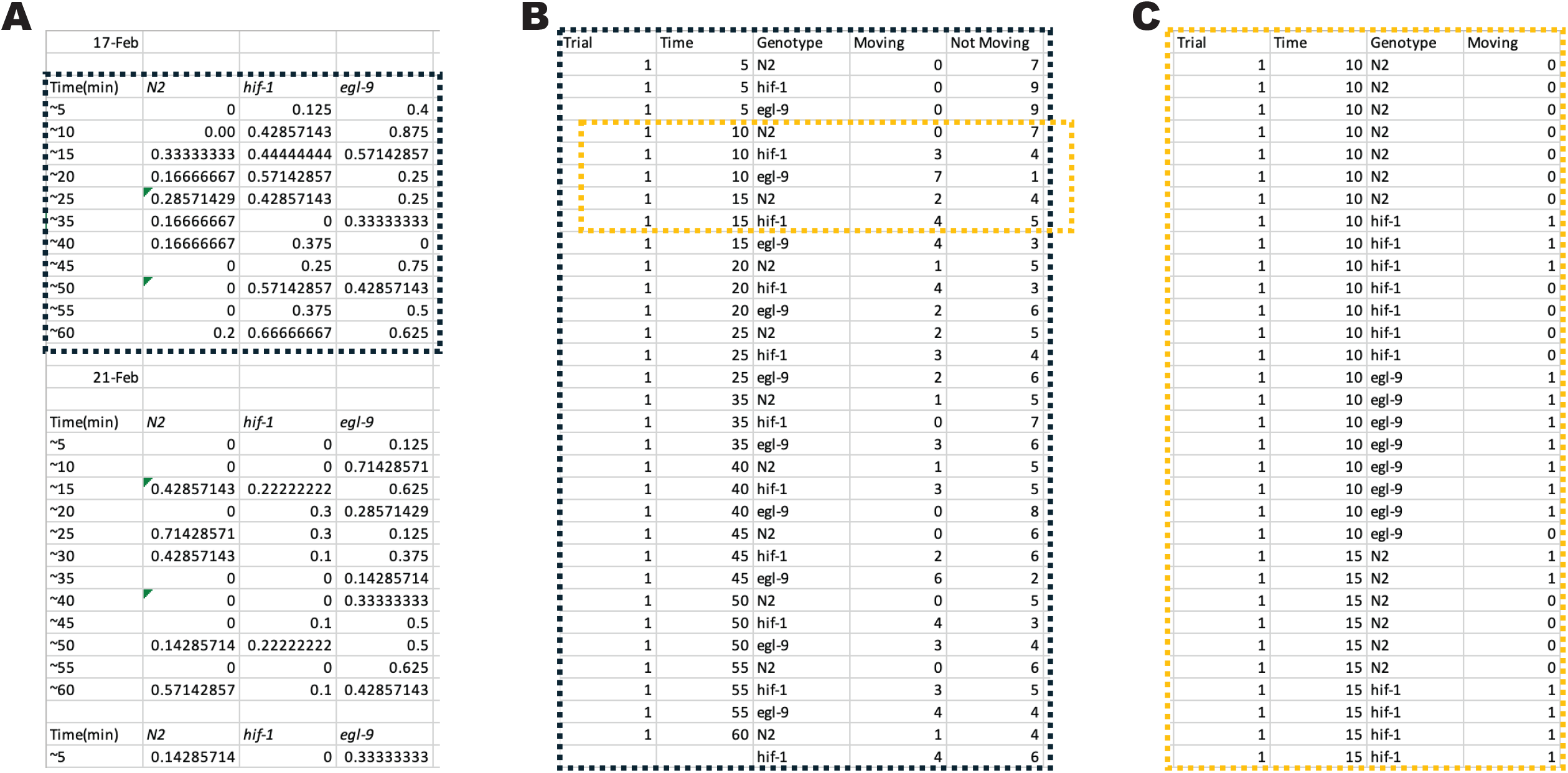
Data Hygiene and Optimization for R import. Datasheet in **A)** shows three trials with information organized into three separate tables. The inset indicated by the dashed box is expanded into an optimized format in **B)** which now has all information organized into columns with that can be called as individual variables. **C)** shows the further expanded dashed box inset in B). In C), variables are still organized in columns, but now each individual animal from the experiment is represented on a single row. Reformatting code in Supplementary information can be used to expand B to C.

### Logistic regression applications for more complex datasets: adding conditions/variables

It is often of interest to include multiple comparisons with the control strain in an experiment. A substantial advantage of logistic regression over CMH is the former can readily accommodate more complex situations. For example, logistic regression accommodates the type of datasets produced when comparing multiple genetic conditions or performing an RNAi screen; in other words, logistic regression accommodates experimental designs with multiple perturbations. In the published example that follows, the authors examined the effect of fertility on exopher rates by monitoring exopher production in hermaphrodites under three conditions: unmated, mated with sterile males, and mated with fertile males^1^. To detect any significant changes across all groups, we included the three conditions within a single spreadsheet as input for our analysis (See: https://github.com/Randrowski/Logistic-Regression-for-Biologists.git). We included code for reformatting/expansion of the data from total instances (i.e., total exophers produced within a trial) to one observation per animal (i.e., did a particular animal produce an exopher, yes/no). This data expansion step generates a spreadsheet optimally formatted for analysis.

Using the reformatted data spreadsheet as input, the analysis code that we provide includes an initial calculation of the mean and standard error of the percentage. We plotted the data to take an initial look at the sample averages and variation between the groups, and then we calculated the logistic regression. This included a determination of whether the data were statistically different (p-value), how different were the means and in which direction was this difference (z-value), and what was the likelihood of the neuron producing an exopher under these conditions (odds ratio). The logistic regression code we generated makes specific comparisons among multiple groups monitored in the same study; here, we consider only three (unmated, mated with sterile males, and mated with fertile males). Even when the numbers of conditions increase, the logistic regression can easily accommodate any number of comparisons. In this case, co-culturing hermaphrodites and sterile males resulted in no significant change in exopher rates compared to unmated hermaphrodites (p=0.756). However, mating to fertile males increased the exopher rate significantly compared to the unmated control (p=0.0000473, and a positive z-score of 4.068, indicating an increased frequency compared to control) (Figure 1C). Our analysis contains code that allows one to specify the control treatment, which should be kept in mind during data interpretation as a positive z-value indicates that the frequency in the experimental group is greater than that of the control, whereas a negative z-value indicates it is less than in the control.

### Logistic regression applications for more complex datasets: assessing how a phenotype can change over time

Often a phenotype is changeable over the timeframe in which it is scored. The final example we consider here is an established anoxia reoxygenation model that involves a predictable behavioral response in *C. elegans*. Anoxia is an environmental condition of extreme oxygen deprivation (<0.1% O_2_). When exposed to anoxia, nematodes enter a reversible hypometabolic state called suspended animation in which movement, feeding, egg laying, and cell division (except anaphase) are arrested^9^. This behavior is reversible, as nematodes reintroduced to normal oxygen levels can resume activity and develop into reproductive adults.

Reoxygenation recovery is genetically variable, and mutants of the Hypoxia Inducible Factor (HIF) pathway can be tested for differences in mobility in response to reoxygenation. Briefly, the HIF pathway includes the transcriptional effector HIF-1, which is activated under conditions of oxygen deprivation, and the prolyl hydroxylase EGL-9, which senses oxygen and inhibits the activity of HIF-1 under normal oxygen conditions^10^. We previously demonstrated that at ten minutes post reoxygenation, *egl-9(sa307)* loss-of-function mutants show an enhanced ability to recover when compared to the wild-type (N2) and *hif-1(ia4)* strains^11^. Although this result is robust (Figure 3A, ten minutes post reoxygenation), the dynamics of reoxygenation observed over longer durations reveal unexpected patterns that may be biologically informative (Figure 3B, one hour post reoxygenation). We anticipated that representation of categorical proportions and trends of each mutant could be rigorously assessed using a logistic regression analysis, which we describe below.

**Figure 3.**
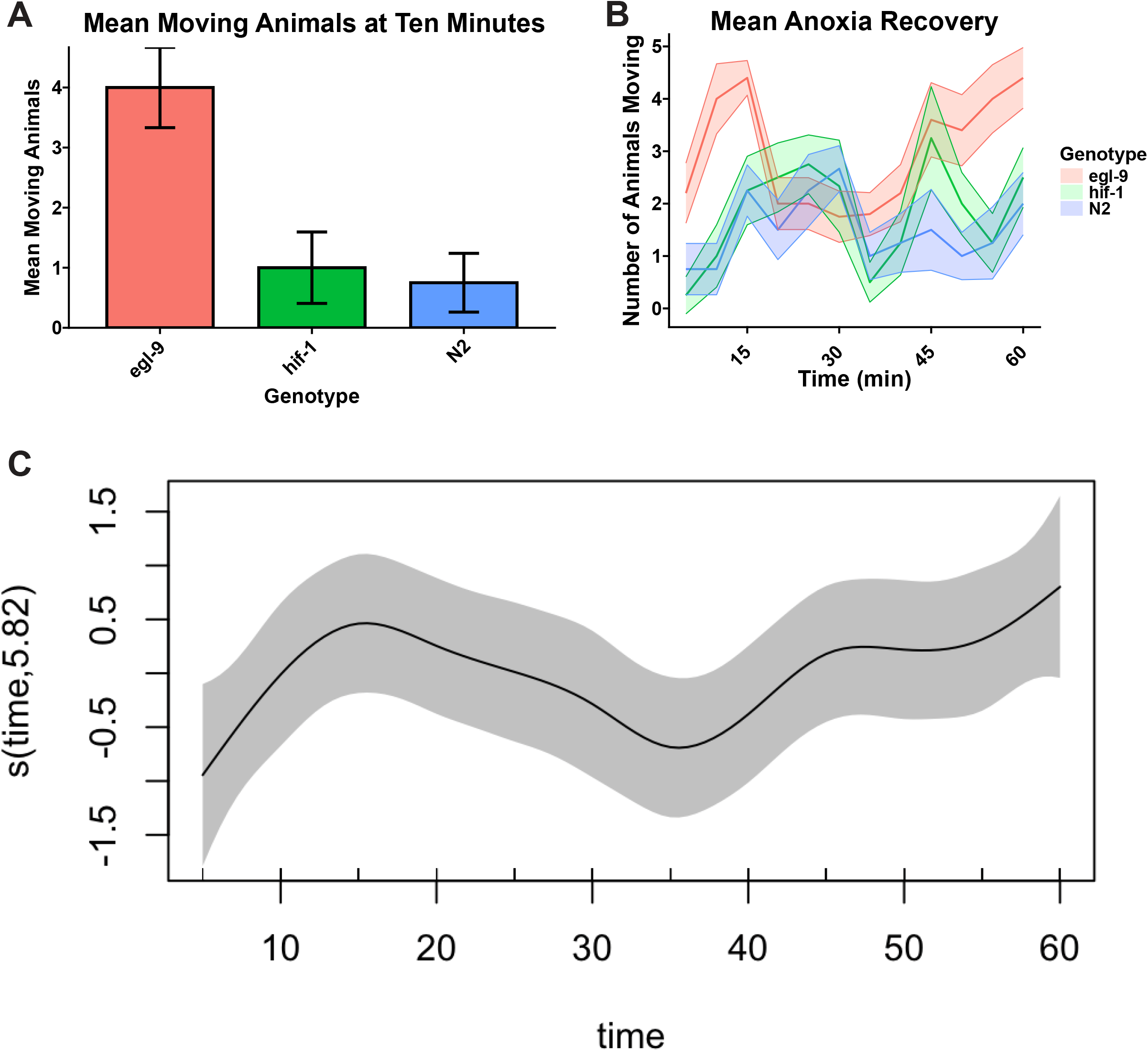
Analysis of anoxia recovery with logistic regression models. **A)** Bar plot showing the average proportion of animals moving at the ten minute time point of the anoxia recovery assay. Bars represent means of trials (shown as points). Error bars represent SEM. ** = p<0.01 **B)** Time course plot of anoxia recovery for all time points. Solid lines represent mean animals moving at each time point. Error is represented by shaded areas as SEM. **C)** A plot showing the generalized additive model (GAM) for all three genotypes across all time points.

### Issues with blanket ANOVA statistics that can better be addressed with logistic regression

In our study, we counted animals that were moving or not moving (categorical data) every five minutes post reoxygenation, initially generating a curve that represents repeated measures along a time course (Figure 3B). We note, however, that the curve that we generated violates the assumptions of many statistical tests like t-tests and ANOVA. The first of these assumptions is the independence of measurements because the same animals are being measured from time point to time point^12^. Second, complications arise with the representation of the outcome as a proportion (a typical way of presenting the data in the field): if four of ten animals are moving, then the proportion of animals moving is 0.4 at a given time point. The plotting of proportion fails to capture the variability of the biology being observed by not considering variability in the denominator of that proportion—the total number of animals scored for that point. In our example, post reoxygenation animals often explore off the agar and either meet premature desiccation death or escape detection at the time of first counting (but make their way back to the moist safety of the agar later) such that the absolute population (i.e., the denominator) effectively changes from time point to time point within a single plate. Logistic regression accounts for a variable denominator by allowing one to consider <u>both outcomes</u>, a constant denominator and a variable denominator (in R, this is accomplished with a cbind function), whereas ANOVA does not account for a variable denominator, as ANOVA relies on a single continuous variable in the form of a proportion^13^. Although there is consideration of overall sample size in ANOVA, the individual data are lost once the data have been collapsed into a proportion. It is important to emphasize that these aspects of data (non-independent sampling; changing absolute population numbers) are frequently present in basic research^13^ and yet the more appropriate logistic regression statistics are not commonly applied.

### Logistic regression program

We chose R, an open source and relatively accessible coding language, to conduct our statistical analyses. The R code is split into three sections, which deal with (1) installation of necessary packages and importation of an excel spreadsheet, (2) visualization of data, and (3) execution of line analysis. We emphasize that three main types of analyses are tested here (CMH, ANOVA, logistic regression), but these only scratch the surface of regressions and adaptations that can be applied. For example, we applied a generalized additive model (GAM) to the behavioral data set to show the additive effect of the reoxygenation behavior across all genotypes (Figure 3C). GAMs are regression models that allow for non-linear relationships between a predictor (such as age or time) and an outcome. Essentially, the model assumes the relationship is a smooth, but unspecified function, rather than a line, and the shape of the curve informs on the data or process assessed. GAMs are appropriate when there are one or more predictors that are unlikely to have a linear relationship with the outcome. The GAM is useful to extrapolate trends from data so that future genotypes can be compared to the general trend to reveal adherence or deviation. Code and details on how to run these analyses are available at github.com (https://github.com/Randrowski/Logistic-Regression-for-Biologists.git).

### Concluding remarks

Biologists are tasked with observing and describing phenomena within nature. As scientists, we must reduce our observations to variables that can be repeatedly observed in many circumstances. Statistical analysis allows us to compare these variables. We describe here the analysis of two biological phenomena: neuronal exopher production (Figure 1) and changes in locomotory movement over time (Figure 3). Since these two phenomena can be reduced to a binary variable (present/absent and yes/no, respectively), logistic regression allows us to implement a rigorous and informative statistical analysis (Table 1). The simplest way to accomplish this analysis is through R coding, which over half of the authors on this publication were not familiar with at the outset of this collaboration. While we provide sample code, R is open source and easily accessible; there are many talented programmers who share their code widely. In addition, the growing utility of AI machine learning models, such as Google’s “Gemini” engine, has proven themselves to be invaluable resources for troubleshooting R code. During the preparation of our datasets for import into R, we realized that even if logistic regression is not a good fit for a data set, improved data hygiene is invariably a benefit to a lab’s reproducibility and analysis efforts (Figure 2). We argue that logistic regression should be a default consideration for analysis of categorical data in biology in the hope that the arguments presented here will facilitate more rigorous analyses within our scientific community.

## References

(1) Wang, G.; Guasp, R.; Salam, S.; Chuang, E.; Morera, A.; Smart, A. J.; Jimenez, D.; Shekhar, S.; Friedman, E.; Melentijevic, I.; et al. Mechanical force of uterine occupation enables large vesicle extrusion from proteostressed maternal neurons. eLife 2024/7/31, 13. DOI: 10.7554/eLife.95443.2.

(2) Spitzer, K.; Pelizzola, M.; Futschik, A. MODIFYING THE CHI-SQUARE AND THE CMH TEST FOR POPULATION GENETIC INFERENCE: ADAPTING TO OVERDISPERSION. Annals of Applied Statistics 2020, 14 (1), 202–220, Article. DOI: 10.1214/19-AOAS1301.

(3) Aggarwal, R.; Ranganathan, P. Common pitfalls in statistical analysis: Linear regression analysis. Perspectives in Clinical Research Apr-Jun 2017, 8 (2). DOI: 10.4103/2229-3485.203040.

(4) Hosmer Jr, D. W.; Lemeshow, S.; Sturdivant, R. X. Applied logistic regression; John Wiley & Sons, 2013.

(5) Melentijevic, I.; Toth, M. L.; Arnold, M. L.; Guasp, R. J.; Harinath, G.; Nguyen, K. C.; Taub, D.; Parker, J. A.; Neri, C.; Gabel, C. V.; et al. C. elegans neurons jettison protein aggregates and mitochondria under neurotoxic stress. Nature 2017, 542 (7641), 367–371. DOI: 10.1038/nature21362.

(6) Arnold, M. L.; Cooper, J.; Grant, B. D.; Driscoll, M. Quantitative Approaches for Scoring in vivo Neuronal Aggregate and Organelle Extrusion in Large Exopher Vesicles in C. elegans. JoVE (Journal of Visualized Experiments) 2020/09/18, (163). DOI: 10.3791/61368.

(7) Hector, A.; Felten, S. V.; Schmid, B. Analysis of variance with unbalanced data: an update for ecology & evolution. Journal of Animal Ecology 2010/03/01, 79 (2). DOI: 10.1111/j.1365-2656.2009.01634.x.

(8) McDonald, J. H. Handbook of biological statistics; sparky house publishing Baltimore, MD, 2009.

(9) Padilla, A. P.; Ladage, L. M. Suspended animation, diapause and quiescence. Cell Cycle 2012, 11 (9), 1672–1679. DOI: 10.4161/cc.19444.

(10) Powell-Coffman, J. A. Hypoxia signaling and resistance in C. elegans. Trends in Endocrinology & Metabolism 2010/07/01, 21 (7). DOI: 10.1016/j.tem.2010.02.006.

(11) P, G.; EC, P.; A, T.; N, S.-V.; C, R. Anoxia-reoxygenation regulates mitochondrial dynamics through the hypoxia response pathway, SKN-1/Nrf, and stomatin-like protein STL-1/SLP-2 - PubMed. PLoS genetics 2013, 9 (12). DOI: 10.1371/journal.pgen.1004063.

(12) Aarts, S.; van den Akker, M.; Winkens, B. The importance of effect sizes. DOI: 10.3109/13814788.2013.818655.

(13) Z, Y.; M, G.; Sf, G.; L, C.; Tc, H.; X, X. Beyond t test and ANOVA: applications of mixed-effects models for more rigorous statistical analysis in neuroscience research - PubMed. Neuron 01/05/2022, 110 (1). DOI: 10.1016/j.neuron.2021.10.030.

